# Oral 4’fluorouridine provides postexposure protection against lethal Nipah virus infection

**DOI:** 10.64898/2026.02.21.707194

**Authors:** Robert W. Cross, Declan D. Pigeaud, Victoriya Borisevich, Krystle N. Agans, Mack B. Harrison, Rachel O’Toole, Abhishek N. Prasad, Thomas W. Geisbert

## Abstract

There are no approved medical countermeasures for combatting Nipah virus (NiV) which causes regular outbreaks in humans and animals in South and Southeast Asia with mortality rates in humans ranging from 40% to more than 90%. Recently, it was shown that 4’-fluorouridine (4’-FlU; EIDD-2749), an orally available ribonucleoside analog, protected guinea pigs and nonhuman primates from lethal challenge with Lassa virus and that 4’-FlU has *in vitro* antiviral activity against NiV. Here, we assessed the postexposure protective efficacy of 4’-FlU in a lethal hamster model of NiV infection. Daily treatment with 4’-FlU beginning 3 days after exposure to NiV resulted in complete protection from lethal infection. Our findings support the further development of 4’-FlU as a therapy for NiV disease.

## Introduction

Nipah virus (NiV) is a member of the genus *Henipavirus* (family *Paramyxoviridae*) that can cause severe respiratory illness and encephalitis in animals and humans is among the deadliest pathogens known to man with case fatality rates greater than 90%^1-3^ in some outbreaks. NiV is a zoonic disease with bats from the *Pteropus* genus, commonly known as flying foxes, serving as the major natural reservoir host although there is evidence of circulation in other pteropid bats^4^. In several animal species and in humans, the primary pathological observation of NiV infection is a severe systemic and often fatal neurologic and/or respiratory disease^5-7^. NiV can also cause relapsing encephalitis that occurs from a few months to more than 10 years after an acute infection. This is due to recrudescence of NiV replication in the central nervous system (CNS)^8-10^. NiV has a broad species tropism and can cause disease in hamsters, guinea pigs, ferrets, cats, dogs, pigs, horses, squirrel monkeys, African green monkeys, and humans^7,11^ and has been shown to infect chicken embryos^12^.

While NiV infection of humans mostly occurs by contact with infected animals (e.g., consumption of raw date palm sap), multiple rounds of person-to-person transmission of the Bangladesh strain of NiV (NiVB) have been reported^13,14^. Outbreaks of fatal NiV disease in countries including the Philippines, Bangladesh, and India have become more common. The current 2026 outbreak of NiV continues the almost yearly occurrence of episodes in the endemic areas since 2001^3,15^. Importantly, there are no approved preventive vaccines or therapeutics for disease caused NiV. Because of the high case fatality rates, lack of licensed medical countermeasures, ability to be transmitted by respiratory droplets, and the potential for deliberate misuse NiV is classified as a biological safety level 4 (BSL-4) virus and Tier 1 priority pathogen by US Government agencies^16^. NiV is included on the World Health Organization’s (WHO) 2018 List of Priority Pathogens^17^ as well as the *Coalition for Epidemic Preparedness Innovations (CEPI)* list of Priority Diseases^18^.

Substantial progress has been made in developing vaccines against NiV with several platforms including a recombinant soluble version of the HeV attachment glycoprotein G (HeVsG) completely protecting AGM in preclinical studies and being tested in phase 1 clinical trials^2,19^. In regard to antivirals, a number of postexposure and therapeutic treatments have been explored as medical countermeasures in preclinical animal challenge models of NiV infection including chloroquine, ribavirin, remdesivir, polyIC12U, and heptad peptide fusion inhibitors but all with minimal to no effectiveness^20,21^. The most promising postexposure intervention to date has been human monoclonal antibodies (mAb). The human mAb m102.4, reactive to the NiV G glycoprotein, completely protected African green monkeys (AGM) against a high dose respiratory tract challenge with the Malaysia strain of NiV (NiVM) when given 5 and 7 days after exposure^22^. However, this same dose regimen of m102.4 did not protect NiVB-infected AGM from lethal disease but did protect against NiVB when given earlier on days 3 and 5 after exposure^23^. m102.4 has similar *in vitro* neutralization activity against both the NiVB and NiVM strains^24^, suggesting that the NiVB isolate is more virulent in AGM which is consistent with the higher case fatality rates reported for NiVB in humans^1-3^. Importantly, m102.4 was assessed in a phase 1 safety trial and was administered under a compassionate use protocol for postexposure prophylaxis after a NiV exposure to one person in the United States and in more than 15 people exposed to the closely related Hendra virus in Australia^25^. More recently we showed that a human mAb against the prefusion conformation of the F glycoprotein, hu1F5, completely protected AGM when given as a single treatment at doses as low as 10 mg/kg on day 5 after exposure to NiV and we demonstrated that hu1F5 is largely resistant to the generation of escape mutations^26^. Hu1F5 is currently in a phase 1 clinical trial^21^.

While results with human mAb-based therapies for NiV infection are encouraging, the mAbs must be administered by slow bolus intravenous (i.v.) infusion which presents logistical challenges and additional costs. Also, conventional mAb therapeutics have limited ability to cross the blood-brain barrier due to their large size and their hydrophilic composition makes treatment of advanced cases of neurotropic infections like NiV difficult to address with mAbs alone. There is a clear need for oral antivirals against henipaviruses. Regarding oral antivirals against NiV, a recent study showed that 4’-fluorouridine (4’-FlU; also known as EIDD-2749) can inhibit NiV *in vitro*^27^. In addition, 4’-FlU protected guinea pigs against challenge with arenaviruses including Lassa virus (LASV) and Junin virus when given late in the disease course^27^ and completely protected AGM against LASV challenge also administered beginning when the animals were severely ill^28^. 4’-FIU is a ribonucleoside analog currently in pre-clinical development that has broad-spectrum activity against several different families of negative-strand and positive-strand RNA viruses. In addition to having activity against arenaviruses^27^, 4’-FIU can inhibit the replication of respiratory syncytial virus (RSV) and other paramyxoviruses^29^, severe acute respiratory syndrome coronavirus 2 (SARS-CoV-2)^29^, and influenza A viruses^30,31^. Recently, 4’-FIU was shown to be efficacious against Venezuelan equine encephalitis virus, another neurotropic virus, suggesting that it can retain potency beyond the blood brain barrier^31^. 4’-FIU inhibits RNA-dependent RNA polymerase (RdRp) function and causes stalling of both RSV and SARS-CoV-2 polymerases^29^ while incorporation of 4’-FIU-triphosphate triggered immediate chain termination of the influenza A virus RdRp complex^31^.

Given that 4’-FIU has *in vitro* activity against NiV and has shown strong *in vivo* activity against other RNA viruses here we assessed the postexposure protective efficacy of 4’-FIU against NiV in a lethal hamster model of NiV disease.

## Results

To confirm previous findings^27^ we first assessed the *in vitro* antiviral activity of 4’-FIU against our isolate of NiVB. Vero 76 monolayers were inoculated with NiVB in the presence of 4’-FIU (100 µM to 0.391 µM), and supernatants were collected 48 hours after infection. Plaque titrations of supernatants were used to generate a dose-response curve from which the 50% inhibitory concentrations (IC) were interpolated for values derived from a control plate infected in the absence of 4’-FIU. 4’-FIU showed strong inhibitory activity against NiVB, with calculated IC50 value of 0.228 µM (**data not shown**).

To determine if 4’-FIU can be effective as a postexposure treatment against NiV exposure, eighteen Syrian golden hamsters were challenged with 10,000 PFU of NiVB by intraperitoneal (i.p.) injection. One group of animals (n = 6) were treated beginning right after NiVB challenge with 5 mg/kg doses of 4’-FIU orally administered once daily for a total of 10 consecutive days while a second group of animals (n = 6) were treated beginning 3 days post infection (DPI) with 5 mg/kg doses of 4’-FIU orally administered once daily for a total of 10 consecutive days. Treatment was withheld from the third group of in-study positive control animals (n = 6). Three hamsters in the positive control group showed a constellation of clinical signs consistent with NiV disease reported in previous studies^32,33^ including ruffled fur, labored breathing, head tilt, tremors, and/or hind limb paralysis and succumbed on 7 DPI (**Figure 1a-d**) The remaining two positive control animals displayed similar signs and succumbed on 10 and 11 DPI, respectively. Only a single animal in the control group exhibited notable weight loss (∼ 10%, **Supplemental Figure 1c**). All 12 treated animals remained healthy until 21 DPI when one animal in the early treatment group abruptly succumbed (**Figure 1a-d**). Two additional animals in the early treatment group developed clinical signs of NiV disease including ruffled fur, labored breathing, head tilt, front limb weakness, and/or seizures and succumbed on 23 DPI. The remaining two hamsters in the early treatment group never showed any clinical signs of NiV disease during the course of the study and were healthy at the predetermined 35 DPI study endpoint. Likewise, all six hamsters in the 3 DPI treatment group never showed any clinical signs of NiV disease during the course of the study and were healthy at the predetermined 35 DPI study endpoint (**Figure 1a-d**). None of the animals in either of the 4’-FIU treatment cohorts exhibited notable weight loss (> 5%, **Supplemental Figure 1a, c**), and only mild increases in body temperature were observed in some animals across all cohorts, although animals that succumbed tended to exhibit hypothermia during the terminal phase of disease (**Supplemental Figure 1b, d, f**). Significant differences in the survival curves were noted between both treated groups compared against each other as well as to the untreated positive control group (Holm-Šídák multiplicity-corrected p ≤ 0.020 for all comparisons; Mantel-Cox logrank test, **Figure 1a**). There was no significant difference in the proportional survival of hamsters with treatment initiated on day of challenge (day 0) versus 3 DPI nor 0 DPI versus untreated positive controls (Hochberg multiplicity-corrected p = 0.121 and p = 0.455, respectively; Fisher’s exact test); however, proportional survival significantly differed between the cohort treated beginning 3 DPI and the untreated control cohort (Hochberg multiplicity-corrected p = 0.007; Fisher’s exact test).

**Figure 1:**
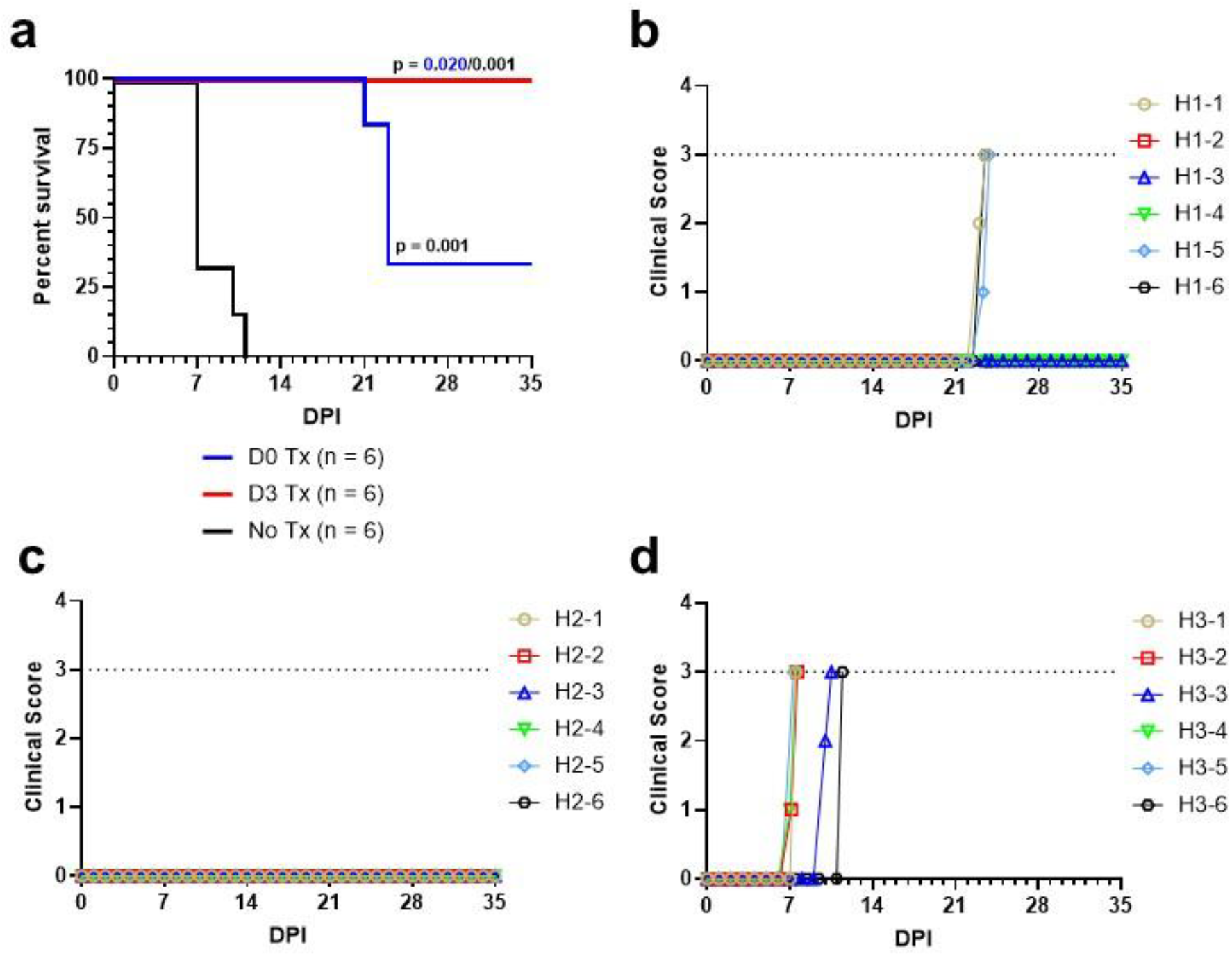
Survival analysis and clinical scoring of hamsters challenged with NiVB and treated with 4’-FIU. **(a)** Kaplan-Meier survival curves for 4’’FIU-treated and untreated (control) hamsters challenged with NiVB. Significance was determined using the Mantel-Cox logrank method with the Holm-Šídák correction for multiple comparisons. Colored p-values denote statistical significance to the same-colored group. All reported p-values are two-tailed. **(b-d)** Clinical scores for hamsters following challenge with NiVB and treatment with 4’-FIU **(a, b)** or no treatment **(d)**. Horizontal dashed lines indicate the score at which humane euthanasia criteria were met.

To determine viral load in mucosal compartments, oral and nasal swabs were collected immediately prior to challenge and at 3, 5 and 7 DPI and assayed for the presence of NiVB viral RNA (vRNA) by RT-qPCR (**Figure 2a-f**). Consistent with NiV hamster i.p. route model only 2/6 hamsters in the positive control group had detectable NiVB vRNA in nasal secretions while 4/6 hamsters in the positive control group had detectable NiVB vRNA in the oral mucosa (**Figure 2e, f**). None of the hamsters from either 4’-FIU-treated group had detectable NiVB vRNA in either nasal or oral secretions (**Figure 2a-d**). Total RNA was extracted from samples of spleen, lung, and brain collected at necropsy from each animal and assayed for the presence of NiVB vRNA by RT-qPCR (**Figure 3a, b**). A significant difference in tissue titer was observed in lung between the 0 DPI FIU-treated group and positive control group and the 3 DPI FIU-treated group and positive control group (**Figure 3a**, p = 0.024 and p = 0.022, respectively; Tukey’s multiple comparisons ANOVA post-hoc test); however, there was a significant difference in titer between animals that survived (n = 8) and animals that succumbed (n = 10) for all tissues (**Figure 3b**).

**Figure 2:**
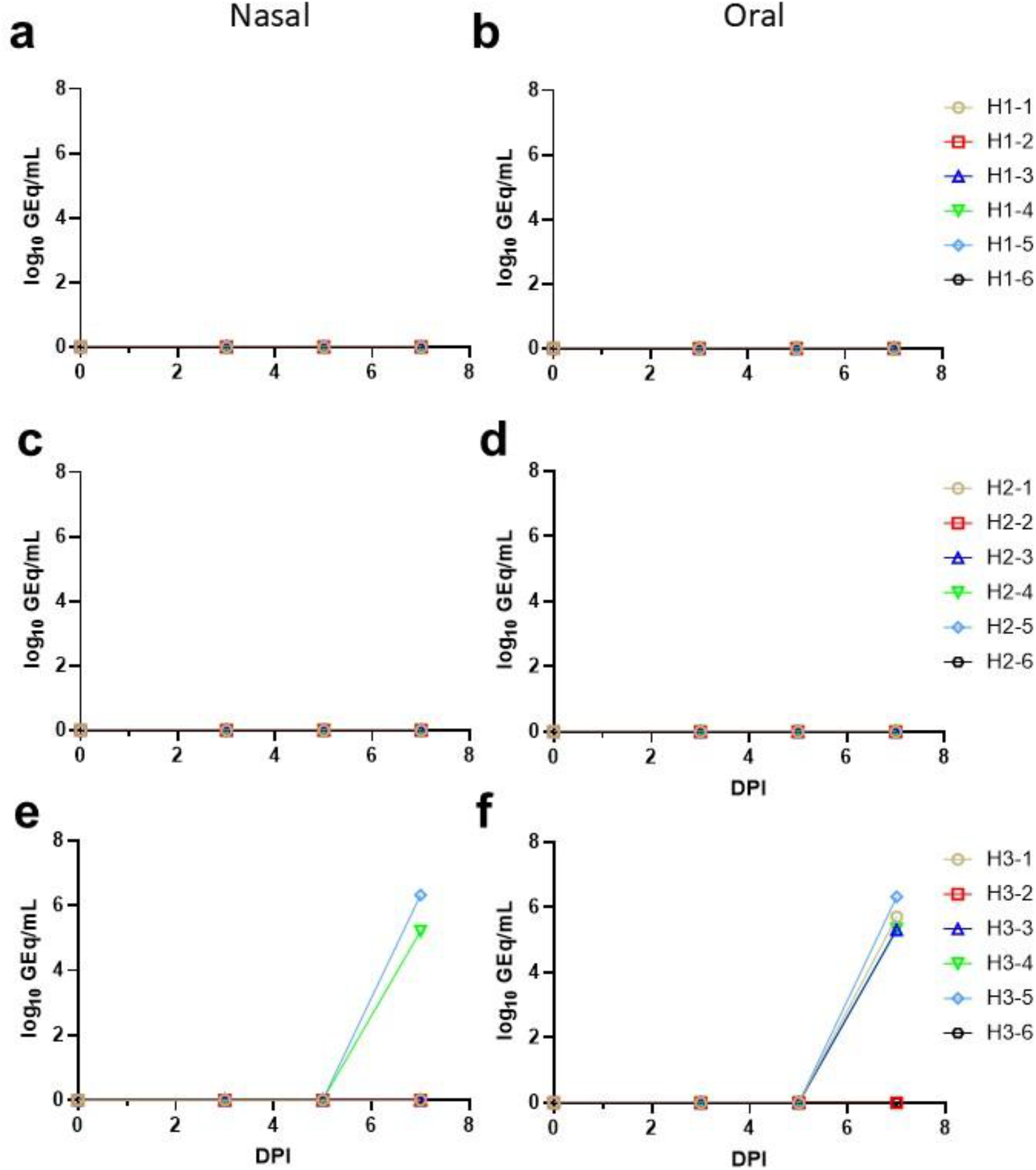
NiVB viral load in nasal and oral secretions. Total RNA was extracted from nasal **(a, c, e)** and oral **(b, d, f)** swabs collected at 0, 3, 5, and 7 DPI and assayed for the presence of NiV vRNA by RT-qPCR. **(a, b)** D0 Tx cohort; **(c, d)** D3 Tx cohort; **(e, f)** No Tx cohort. For all panels, data shown represents the mean genome equivalent titer derived from two replicate assays.

**Figure 3:**
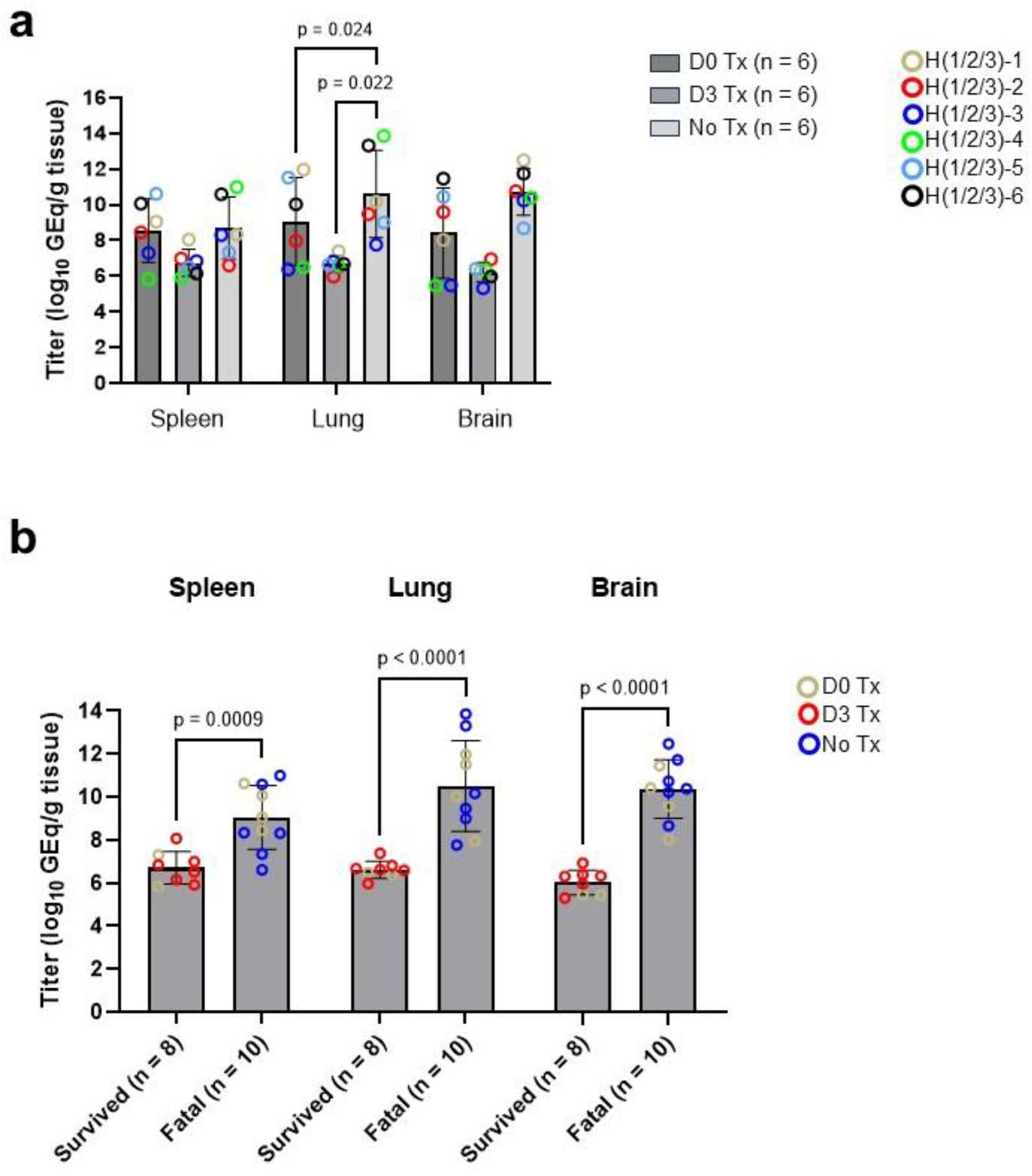
Detection of NiVB vRNA in selected tissues. The indicated tissues were harvested at necropsy and assayed for the presence of NiV vRNA by RT-qPCR. Statistical analysis was performed by comparing NiVB vRNA abundance by cohort **(a)** or by clinical outcome **(b)**. Statistical analysis was performed using 2-way ANOVA with Tukey’s post-hoc test for multiple comparisons (a) or Mann-Whitney U-test (b). Data points shown are the mean values of two replicate assays. All reported values are two-tailed.

## Discussion

The current outbreak of NiVB in India emphasizes the constant threat that NiV poses regionally in South and Southeast Asia as well as globally. Oral antivirals that can confer protection against NiV disease have clear advantages in quickly responding to and containing outbreaks, and therefore their development is essential for future outbreak readiness. The ease of supply, storage, distribution, and in particular the administration of oral antivirals compared to injection or infusion are important considerations and benefits to their use. Additionally, broad spectrum antivirals that are effective across genetically diverse families and strains or variants of viruses are needed to lessen the possibility of resistance mutations that may occur from the employment of more narrowly targeted mAbs or preventive vaccines. Here, we show that the antiviral ribonucleoside analog 4’-FIU protects hamsters from lethal NiVB disease when treatment was started at 3 DPI in a model where untreated positive controls begin to succumb at 7 DPI, with all treated animals surviving to the study endpoint (6/6, 100%).

It is interesting that delaying the initiation of 4’-FIU treatment of hamsters until 3 DPI after exposure to NiVB resulted in substantially greater and complete survival versus the delayed death, but relatively low protection (2/6, 33%) observed in hamsters treated shortly after exposure to NiVB. It is well understood in human medicine that pharmacokinetics (PK) of many compounds can be altered in the context of inflammation and/or infection. In part this may be due to changes in drug absorption, clearance, and tissue distribution caused by inflammatory processes and altered metabolisms within affected tissues, and systemically. As such it is reasonable to assume that similar phenomenon may exist in the context of animal models of disease^34^. Another view to consider is whether there is a critical threshold of viral exposure necessary to mount a protective immune response once the treatment concentration falls below a protective threshold after treatment is stopped. It is possible treating too early with a high enough dose of a drug may curb virus replication and viral antigen exposure and thereby limit or delay development of protective immunity.

Our results support the further development of 4’-FIU for the postexposure prophylaxis and therapeutic treatment of NiV infections. Future studies in the robust AGM model, which is the animal model that most accurately recapitulates human NiV disease^7,11,35^, are needed to further advance the development of 4’-FIU. Along with the use of efficacious preventive vaccines, oral antivirals such as 4’-FIU offer potential to further improve the management of future NiV outbreaks.

## Methods

### Virus and 4’-FlU

The isolate of NiVB used in the study was 200401066, which was obtained from a fatal human case during the outbreak in Rajbari, Bangladesh in 2004 and passaged on Vero-E6 cells three times (GenBank accession number AY988601.1). The cell supernatants were stored at -80°C as ∼ 1 ml aliquots. No detectable mycoplasma or endotoxin levels were measured (< 0.5 EU/ml). 4**’**-fluorouridine (4**’**-FlU; EIDD-2749) was obtained from MedChemExpress (Monmouth Junction, NJ, #HY-146246) and prepared in DMSO as 60 mM working stocks and frozen at -80°C until use.

### Antiviral activity assay

Antiviral activity of 4’-FIU against NiVB was assessed by inoculating monolayers of Vero 76 cells (ATCC CRL-1587) were inoculated with a MOI of 0.01 of NiVB in the presence of two-fold serial dilutions of 4’-FIU (100 µM to 0.391 µM), and supernatants were collected 48 hours after infection. Plaque titrations of supernatants were used to generate a dose-response curve from which the 50% and 90% inhibitory concentrations (IC) were interpolated for values derived from a control plate infected in the absence of 4’-FIU.

### Ethics approval

Animal studies were performed in BSL-4 biocontainment at UTMB in the Galveston National Laboratory and approved by the UTMB Institutional Biosafety Committee and IACUC. Animal research was conducted in compliance with the UTMB IACUC, the Animal Welfare Act, and other federal statutes and regulations relating to animals. The UTMB animal research facility is fully accredited by the Association for Assessment and Accreditation of Laboratory Animal Care and adheres to principles specified in the eighth edition of the Guide for the Care and Use of Laboratory Animals, National Research Council.

### 4’-FlU treatment in hamsters

Eighteen 3 to 5 week-old Syrian golden hamsters were randomized by Excel into two experimental groups of six animals each and one positive control group of six animals. All eighteen hamsters were inoculated with 10,000 PFU of NiV (Bangladesh strain, passage 3) by the intraperitoneal route. Shortly after challenge, all six animals in one experimental group were treated with 5 mg/kg of 4’-FlU (as a 1:10 suspension of 4’FIU in DMSO to 1% methyl cellulose in water vehicle) by oral gavage. These six animals in the first experiment group received daily doses of 4-’FIU for 10 days (0-9 DPI). Beginning on 3 DPI the six animals in the second experiment group were treated with 5 mg/kg of 4’-FlU by oral gavage. These six animals in the second experimental group also received daily doses of 4-’FIU for 10 days (3-12 DPI). The six hamsters in the positive control group were not treated. All eighteen animals were monitored for 35 days for changes in weight, temperature, and clinical appearance. Animals were humanely euthanized at the experimental study endpoint.

### RNA isolation from NiVB-infected hamsters

On procedure days, 100 µl of blood from K2-EDTA collection tubes was collected before centrifugation and added to 600 µl of AVL viral lysis buffer with 6 µl of carrier RNA (Qiagen) for RNA extraction. For tissues, approximately 100 mg was stored in 1 ml of RNAlater (Qiagen) for a minimum of 24 hours. RNA was isolated from tissues after homogenization in 600 µl of RLT buffer and 1% betamercaptoethanol (Qiagen) in a 2-ml cryovial using a tissue lyser (Qiagen) and 0.2-mm ceramic beads. The tissues sampled included lung, spleen, and brain. All blood samples were inactivated in AVL viral lysis buffer, and tissue samples were homogenized and inactivated in RLT buffer before removal from the BSL-4 laboratory. Subsequently, RNA was isolated from blood and swabs using the QIAamp viral RNA kit (Qiagen) and from tissues using the RNeasy minikit (Qiagen) according to the manufacturer’s instructions supplied with each kit.

### Detection of NiVB load

RNA isolated from blood or tissues was assessed using primers and a probe targeting the middle of the *N* gene of NiVB by qRT-PCR with the probe used being 6FAM-5′-CTGCAGGAGGTGTGCTCACGGAGG-3′-TAMRA (Life Technologies) as described previously^26^. NiVB RNA was detected using the CFX96 detection system (Bio-Rad) with one-step probe qRT-PCR kits (Qiagen) using the following cycle conditions: 50°C for 10 minutes, 95°C for 10 seconds, and 45 cycles of 95°C for 10 seconds and 57°C for 30 seconds. Threshold cycle (*C*T) values representing NiVB genomes were analyzed with CFX Manager software, and data are presented as gene expression quantitation (GEq) values. To generate the GEq standard curve, RNA from NiVB challenge stocks was extracted, and the number of genomes was calculated using Avogadro’s number and the molecular weight of the NiVB genome.

## Acknowledgments

The authors wish to thank the UTMB Animal Resource Center for husbandry support of laboratory animals and Drs. Jacquelyn Turcinovic and Zachary Schiffman for assistance with animal studies. This study was supported by funds from the UTMB Department of Microbiology and Immunology to TWG and UC7AI094660 for BSL-4 operations support of the Galveston National Laboratory. Opinions, interpretations, conclusions, and recommendations are those of the authors and are not necessarily endorsed by the University of Texas Medical Branch.

## Author contributions

RWC and TWG conceived and designed the experiments. RWC and VB performed the challenges and treatments. RWC, DDP, VB, MBH, and RO performed procedures and conducted clinical observations. KNA performed the PCR assays. All authors analyzed the data. TWG wrote the paper with additions from RWC and ANP. DDP edited the paper. ANP prepared the Figures. All authors had access to the data and approved the final version of the manuscript.

## References

1. Hossain MJ, Gurley ES, Montgomery JM, Bell M, Carroll DS, Hsu VP, et al. Clinical presentation of nipah virus infection in Bangladesh. Clin Infect Dis. 2008 Apr 1;46(7):977–84.

2. Spengler JR, Lo MK, Welch SR, Spiropoulou CF. Henipaviruses: epidemiology, ecology, disease, and the development of vaccines and therapeutics. Clin Microbiol Rev. 2025 Mar 13;38(1):e0012823.

3. Yadav PD, Baid K, Patil DY, Shirin T, Rahman MZ, Peel AJ, et al. A One Health approach to understanding and managing Nipah virus outbreaks. Nat Microbiol. 2025 Jun;10(6):1272–1281.

4. Clayton BA, Wang LF, Marsh GA. Henipaviruses: an updated review focusing on the pteropid reservoir and features of transmission. Zoonoses Public Health. 2013 Feb;60(1):69–83.

5. Wong KT, Ong KC. Pathology of acute henipavirus infection in humans and animals. Patholog Res Int. 2011;2011:567248.

6. Abdullah S, Tan CT. Henipavirus encephalitis. Handb Clin Neurol. 2014;123:663–70.

7. Pigeaud DD, Geisbert TW, Woolsey C. Animal Models for Henipavirus Research. Viruses. 2023 Sep 22;15(10):1980.

8. Sejvar JJ, Hossain J, Saha SK, Gurley ES, Banu S, Hamadani JD, et al. Long-term neurological and functional outcome in Nipah virus infection. Ann Neurol. 2007 Sep;62(3):235–42.

9. Siva SR, Chong HT, Tan CT. Ten year clinical and serological outcomes of Nipah virus infection. Neurol Asia 2009; 14:53–58

10. Wong KT, Tan CT. Clinical and pathological manifestations of human henipavirus infection. Curr Top Microbiol Immunol. 2012;359:95–104.

11. Geisbert TW, Feldmann H, Broder CC. Animal challenge models of henipavirus infection and pathogenesis. Curr Top Microbiol Immunol. 2012;359:153–77.

12. Tanimura N, Imada T, Kashiwazaki Y, Sharifah SH. Distribution of viral antigens and development of lesions in chicken embryos inoculated with nipah virus. J Comp Pathol. 2006 Aug-Oct;135(2-3):74–82.

13. Nikolay B, Salje H, Hossain MJ, Khan AKMD, Sazzad HMS, Rahman M, et al. Transmission of Nipah Virus - 14 Years of Investigations in Bangladesh. N Engl J Med. 2019 May 9;380(19):1804–1814.

14. Hegde ST, Lee KH, Styczynski A, Jones FK, Gomes I, Das P, Gurley ES. Potential for Person-to-Person Transmission of Henipaviruses: A Systematic Review of the Literature. J Infect Dis. 2024 Mar 14;229(3):733–742.

15. World Health Organization. Nipah virus infection-India. Available at: https://www.who.int/emergencies/disease-outbreak-news/item/2026-DON593 (Accessed February 20, 2026).

16. Federal Select Agent Program. Select Agents and Toxins List. Available at: https://www.selectagents.gov/sat/list.htm (Accessed February 20, 2026).

17. World Health Organization. Prioritizing diseases for research and development in emergency contexts. Available at: https://www.who.int/activities/prioritizing-diseases-for-research-and-development-in-emergency-contexts (Accessed February 20, 2026).

18. Coalition for Epidemic Preparedness Innovations (CEPI). Priority pathogens. Available at: https://cepi.net/priority-pathogens (Accessed February 20, 2026).

19. Lv C, He J, Zhang Q, Wang T. Vaccines and Animal Models of Nipah Virus: Current Situation and Future Prospects. Vaccines (Basel). 2025 Jun 4;13(6):608.

20. Broder CC. Henipavirus outbreaks to antivirals: the current status of potential therapeutics. Curr Opin Virol. 2012 Apr;2(2):176–87.

21. Chan XHS, Haeusler IL, Choy BJK, Hassan MZ, Takata J, Hurst TP, et al. Therapeutics for Nipah virus disease: a systematic review to support prioritisation of drug candidates for clinical trials. Lancet Microbe. 2025 May;6(5):101002.

22. Geisbert TW, Mire CE, Geisbert JB, Chan YP, Agans KN, Feldmann F, et al. Therapeutic treatment of Nipah virus infection in nonhuman primates with a neutralizing human monoclonal antibody. Sci Transl Med. 2014 Jun 25;6(242):242ra82.

23. Mire CE, Satterfield BA, Geisbert JB, Agans KN, Borisevich V, Yan L, et al. Pathogenic Differences between Nipah Virus Bangladesh and Malaysia Strains in Primates: Implications for Antibody Therapy. Sci Rep. 2016 Aug 3;6:30916.

24. Dong J, Cross RW, Doyle MP, Kose N, Mousa JJ, Annand EJ, et al. Potent Henipavirus Neutralization by Antibodies Recognizing Diverse Sites on Hendra and Nipah Virus Receptor Binding Protein. Cell. 2020 Dec 10;183(6):1536-1550.e17.

25. Playford EG, Munro T, Mahler SM, Elliott S, Gerometta M, Hoger KL, et al. Safety, tolerability, pharmacokinetics, and immunogenicity of a human monoclonal antibody targeting the G glycoprotein of henipaviruses in healthy adults: a first-in-human, randomised, controlled, phase 1 study. Lancet Infect Dis. 2020 Apr;20(4):445–454.

26. Zeitlin L, Cross RW, Woolsey C, West BR, Borisevich V, Agans KN, et al. Therapeutic administration of a cross-reactive mAb targeting the fusion glycoprotein of Nipah virus protects nonhuman primates. Sci Transl Med. 2024 Apr 3;16(741):eadl2055.

27. Welch SR, Spengler JR, Westover JB, Bailey KW, Davies KA, Aida-Ficken V, et al. Delayed low-dose oral administration of 4’-fluorouridine inhibits pathogenic arenaviruses in animal models of lethal disease. Sci Transl Med. 2024 Nov 20;16(774):eado7034.

28. Cross RW, Turcinovic J, Prasad AN, Borisevich V, Agans KN, Deer DJ, et al. Oral 4’-fluorouridine rescues nonhuman primates from advanced Lassa fever. Nature. 2026 Jan 7. doi: 10.1038/s41586-025-09906-y.

29. Sourimant J, Lieber CM, Aggarwal M, Cox RM, Wolf JD, Yoon JJ, et al. 4’-Fluorouridine is an oral antiviral that blocks respiratory syncytial virus and SARS-CoV-2 replication. Science. 2022 Jan 14;375(6577):161–167.

30. Lieber CM, Plemper RK. 4’-Fluorouridine Is a Broad-Spectrum Orally Available First-Line Antiviral That May Improve Pandemic Preparedness. DNA Cell Biol. 2022 Aug;41(8):699–704.

31. Lieber CM, Aggarwal M, Yoon JJ, Cox RM, Kang HJ, Sourimant J, et al. 4’-Fluorouridine mitigates lethal infection with pandemic human and highly pathogenic avian influenza viruses. PLoS Pathog. 2023 Apr 17;19(4):e1011342.

32. Wong KT, Grosjean I, Brisson C, Blanquier B, Fevre-Montange M, Bernard A, et al. A golden hamster model for human acute Nipah virus infection. Am J Pathol. 2003 Nov;163(5):2127–37.

33. Pigeaud DD, Borisevich V, Agans KN, Harrison MB, O’Toole R, Martinez J, et al. A single-cycle recombinant VSV vaccine displaying the Hendra virus glycoprotein uniformly protects against Hendra and Nipah virus challenge. A single-cycle recombinant VSV vaccine displaying the Hendra virus glycoprotein uniformly protects against Hendra and Nipah virus challenge. NPJ Vaccines. 2026 Jan 3;11(1):21.

34. Martinez MN, Greene J, Kenna L, Kissell L, Kuhn M. The Impact of Infection and Inflammation on Drug Metabolism, Active Transport, and Systemic Drug Concentrations in Veterinary Species. Drug Metab Dispos. 2020 Aug;48(8):631–644.

35. Geisbert TW, Daddario-DiCaprio KM, Hickey AC, Smith MA, Chan YP, Wang LF, et al. Development of an acute and highly pathogenic nonhuman primate model of Nipah virus infection. PLoS One. 2010 May 18;5(5):e10690.

