## Supplementary Figure 1 for "Oral 4’fluorouridine provides postexposure protection against lethal Nipah virus infection" (from another version of this preprint: version 1, 2026-02-22)

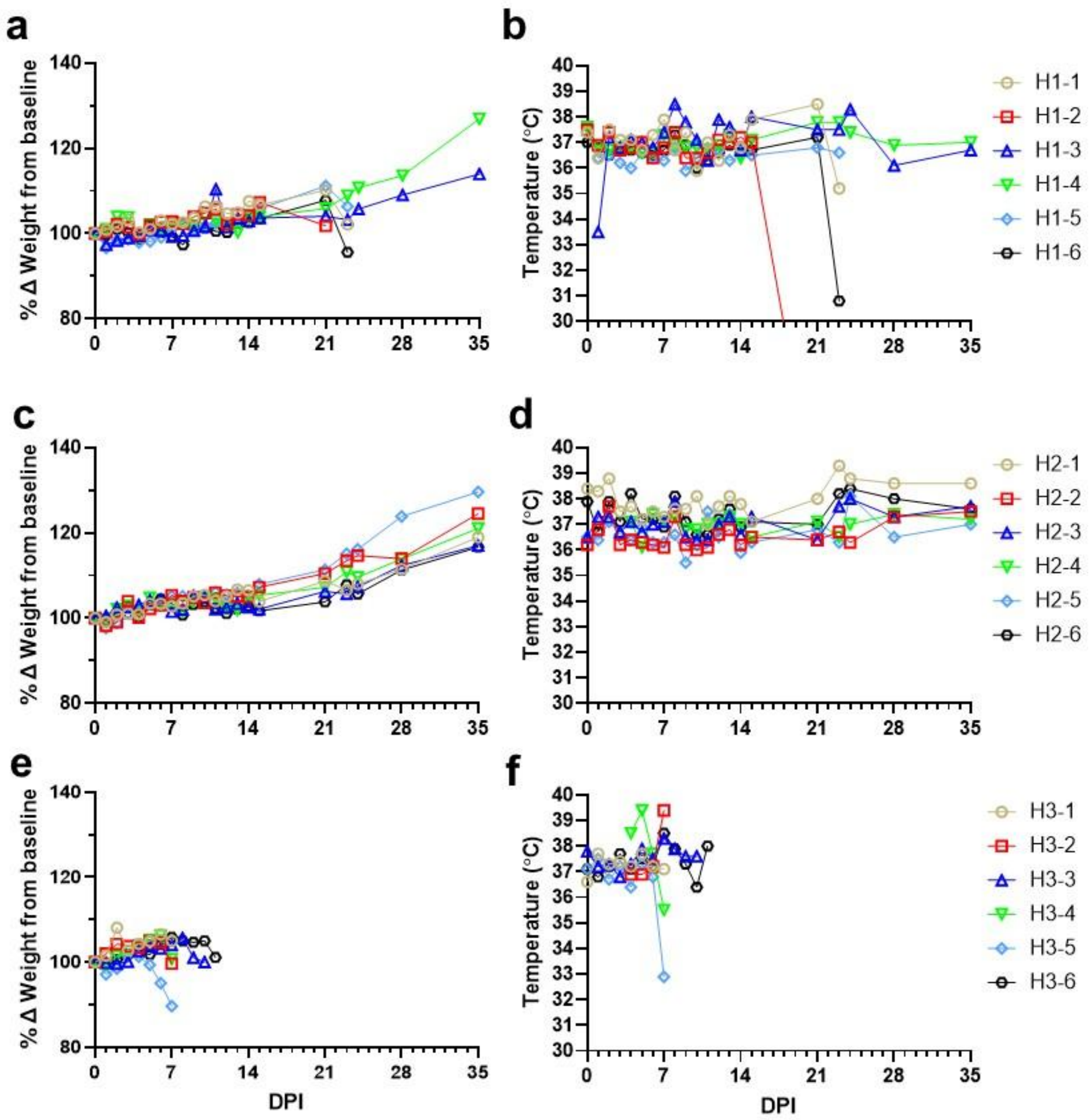


**Supplementary Figure 1: Weight and temperature monitoring of hamsters following challenge with NiVB and treatment with 4`-FIU.** Weight was measured at the indicated timepoints **(a, c, e)**. Temperature was monitored using telemetry and scanned at the indicated timepoints **(b, d, f)**.
